# Is *APOE ε2* always a protective allele? Deviations in Hardy-Weinberg equilibrium in admixed Brazilian elderly individuals

**DOI:** 10.64898/2026.06.20.733520

**Authors:** GN Santos, PH Sebe, Carlos, MALS Alexandria, LA Veronezz, IB Ignácio, SLG Paço, Diogo Meyer team, G Bandeira, MU Bardella, Claudia Kimie Suemoto, Renata Elaine Paraizo Leite, MS Naslavsky

## Abstract

The APOE gene is a critical determinant of human healthspan and longevity, with the rare *ε2* allele traditionally viewed as a universal protective factor against Alzheimer s disease (AD) and a driver of exceptional lifespan. However, this protective paradigm is predominantly derived from European-centric cohorts, leaving the evolutionary and clinical impacts of *ε2* across diverse, highly admixed populations largely unknown due to a lack of local ancestry (LA) resolution. To investigate how local genomic backgrounds modulate *APOE* survival dynamics we analyzed two Brazilian sample collection of older adults from São Paulo city: the Biobank for Aging Studies (BAS, n = 716), a post-mortem autopsy study of naturally deceased individuals; and the Health, Well-being and Aging Study (SABE, n = 952), a census-based elderly sample collection. We evaluated deviations from Hardy-Weinberg equilibrium (HWE) using robust permutation-based models to capture ongoing selective and mortality pressures at the *APOE* locus. While global *APOE* frequencies adhered to HWE, integrating LA unveiled striking, mirrored ancestral deviations. Our findings reveal that *APOE ε2* homozygotes with African ancestry significantly contribute to deviations from HWE in the BAS, with an excess of ε2AFR/ε2AFR homozygotes observed (p = 0.0196). These distinct HWE deviations demonstrate that an African LA background acts as a genetic buffer, attenuating the phenotypic extreme effects of *APOE* alleles. Furthermore, we observed an excess of the *ε4* European haplotypes in the BAS, which is consistent with a mortality pressure allelic effect in the European LA context. Conversely, the *ε4*AFR/*ε4*AFR combination was overrepresented in the SABE. While this buffering mechanism mitigates *ε4* toxicity, it simultaneously dampens the exceptional longevity advantage typically conferred by the *ε2* allele, leading to its neutral accumulation in the post-mortem cohort. Our study challenges the “one-size-fits-all” assumption of *APOE* biomarkers, demonstrating that *ε2* protective mechanisms are context-dependent and modulated by local genomic backgrounds in admixed populations.

## INTRODUCTION

The *APOE* gene is a critical determinant of human healthspan, with its polymorphic variants exhibiting opposing effects on neurodegeneration and aging [1–10]. Specifically, the *APOE ε2* allele is widely regarded as a protective factor against Alzheimer’s disease (AD) and has been robustly associated with longevity [1, 7, 11–14]. Conversely, the *APOE ε4* allele represents the strongest genetic risk factor for late-onset AD, with its attributable risk ranging between 40–65%, depending on *ε4* dosage (homozygotes vs. heterozygotes) and the sociodemographic characteristics of the study population [4, 10, 14–16].

The *APOE ε2* allele is the rarest among *APOE* common alleles (*ε3* and *ε4)*, with a global frequency of less than 10%, although it is highly variable across distinct populations [1, 14, 17–19]. This allele is also a replicable variant in GWAS for longevity. Its protective effects associated with reduction of amyloid and Tau accumulation, could confound the *APOE ε2* effects on longevity. Because *APOE ε2* and *APOE ε4*, respectively, reduce and increase the risk of AD, there is a critical need to address whether the effects of *APOE* genotype on longevity are mediated through such AD risks [6, 13, 20]. Complementary evidence already showed that the protective effect of *APOE ε2* allele is independent of AD, although its mechanisms are still poorly understood [1, 2, 12, 14].

The *APOE* gene is most studied in the context of DA pathology due to the high risk carried by the ε4 allele[21–27]. While it is a well-established genetic factor, the risk associated with this allele may not be uniform across different populations worldwide, as shown by recent studies revealing ancestry-specific modulation of *APOE* effects in AD. Blue *et al*. (2019) showed 39% lower odds of AD in Caribbean Hispanics with African ancestry compared to European ancestry for the *APOE ε4* allele [28]. Rajabli *et al*. (2018) identified a lower risk *APOE ε4* on an African local ancestry (LA) background in Puerto Rican and African American populations [23]. Although AD odds vary among studies, the results consistently show that African ancestry is associated with lower accumulation of AD neuropathology, particularly neuritic plaques (NP). Contributing to the ancestry-driven variability conversation, Schlesinger *et al.* (2013) conducted an analysis in a São Paulo’s admixed population sample, derived from the Brain Aging Study (BAS), and showed that the presence of African Global ancestry (GA) was associated with lower prevalence of NP [24]. Naslavsky *et al*. (2023) further investigated the same sample collection and revealed the higher prevalence of risk in European LA *APOE ε4* carriers in the same population. They also found that increased proportions of African ancestry correlated with worse cognitive performance for individuals with NP severe burden, particularly in *APOE ε4 non*carriers [29].

Due to the low global frequency of *APOE ε2* (<10%) and its underrepresentation in non-European biobanks, previous studies have had a limited understanding of its protective mechanisms across different LA backgrounds, especially under admixed configurations[11–13, 17–19, 30]. Consequently, the protective effects of *APOE ε2* in neurological condition risks have not always been conclusive [6]. Suri *et al*. (2013) addressed this issue, pinpointing most of the recent reports on the protective effect of this allele, described by them as “the forgotten *APOE* allele”. Most reported studies on this subject combine *APOE ε2* heterozygous (often excluding individuals with *APOE ε2*/*ε4* genotypes) or pool the rare homozygous subjects with heterozygotes due to the extremely low frequency of the *ε2* /*ε2* genotype [31]. The challenge also extends to quantify the longevity effect that *APOE ε2* may coffer, given its heterogeneous frequency distribution (eg. 2,9% for mexicans and 8,6% for dominicans), and the fact that these genetic determinants are dynamic and dependent on the environmental history of each population [6, 20].

To advance the understanding of *APOE ε2*’s protective effect on longevity, in the context of LA, we utilized two Brazilian samples: 1) the Biobank for Aging Studies (BAS), hosting samples from a community-dwelling postmortem biobank of naturally deceased elderly from São Paulo city [32–35], and 2) the Health, Well-being and Aging study (SABE), a census-based collection of living older adults with longitudinal health data also from São Paulo city [36, 37]. By leveraging these two sample collections, we were able to analyze and compare *APOE* allelic and genotypic frequencies alongside LA inferences. Crucially, this comparison of the genotype frequencies between a deceased (BAS) and a living (SABE) sample collections allows us to investigate potential survival or longevity effects conferred by the *APOE ε2* allele in Brazilian admixed individuals.

To test for proxies of differential survival or selection pressure across ancestries, we used the Hardy–Weinberg equilibrium (HWE), a foundational principle in population genetics that predicts the stability of allele and genotype frequencies in a population over generations, assuming no evolutionary forces are acting. It serves as a null model to detect evolutionary changes or genetic anomalies, in addition to detecting genotyping artefacts, such as allelic dropout [38]. From a clinical perspective, HWE may act as a tool for estimating genetic risk and understanding disease epidemiology [39]. HWE deviations may signal unique genetic dynamics shaped by ancestry-specific evolutionary pressures [6, 40]. Therefore, we aimed to investigate whether the longevity protective effects of *APOE ε2* in the São Paulo population are modified by or depend on ancestry. Specifically, we assessed whether this dependence could be detected through HWE deviations when comparing the BAS (deceased) and SABE (living) sample cpllections.

## METHODS

### Study Populations and Sampling

Two population-based samples from the city of São Paulo (Brazil) were used in this study. The central goal was to compare a survival sample collection (SABE) against a mortality study (BAS) to investigate the differential effects of *APOE* and LA on mortality risk. The first sample is derived from the SABE (Health, Well-being and Aging), an established census-based longitudinal dataset of older adults (≥ 60 years) collected using a multistage stratified sampling method [36]. The second sample is derived from the Biobank for Aging Studies (BAS), which includes biological samples collected from individuals who died at age 18 or older, sourced from the Sao Paulo Autopsy Service (providing compulsory postmortem verification for deaths due to non-traumatic causes). As a population-based biobank, BAS minimises selection biases inherent to disease-specific or hospital-based autopsy studies. The datasets analysed here received prior approval from the respective local ethics committees and have been previously published [36, 36].

Both samples comprise individuals with various comorbidities, including cardiovascular and neurodegenerative diseases, with prevalence compatible with the age span. Despite notable differences in sampling methods, the BAS is specifically characterized by a high prevalence of vascular endophenotypes (main cause of death is cardiovascular diseases), making it highly relevant for testing the effects of *APOE* on mortality, given its role in cardiovascular disease [32].

Participants (or otherwise legally appointed representatives) of SABE and family members (or otherwise legally appointed representatives) of BAS signed approved consent forms for participation in these studies. Approval followed Brazilian regulation. SABE participants were approved by COEP/CEP/CONEP (Brazilian local and national ethical committee boards) under the following protocols: COEP FSP USP OF.COEP/23/10, CONEP 2044/2014, CEP HIAE 1263-10. BAS approvals:

### Genomic DNA Extraction and *APOE* Genotyping

Genomic DNA was extracted from peripheral blood (SABE) or peripheral blood (BAS) and somatic tissues (frozen brain tissue for BAS, when DNA quality from blood did not reach the minimum quality/quantity measured on Nanodrop) [29, 36]. *APOE* genotyping of common alleles (composed by rs7412 and rs429358 haplotypes) was performed using allele-specific real-time PCR amplification [41] and confirmed by whole-genome sequencing (on SABE, Naslavsky et al., 2022) [36]. Global and local ancestry used genotypes from microarray (Illumina OmniExpress 700k microarray) or whole-genome sequencing for BAS and SABE, respectively. For this study, the primary exposure of interest is the *APOE* genotype. We included all participants with available *APOE* allele-specific genotypes from both collections, resulting in 716 samples from BAS and 952 from SABE. *APOE* common alleles were preferably genotyped directly using allele-specific amplification or after imputation of rs429358 to compose haplotypes. Individuals from both sample collections were classified as carriers of *APOE* genotypes ε2/ε2, ε2/ε3, ε2/ε4, ε3/ε3, ε3/ε4, and ε4/ε4.

### Local Ancestry Inference

LA inference was performed on a subset of 716 individuals with available microarray data from the BAS and a subset of 952 individuals with available LA inference from the whole genome sequencing data from the SABE. Populations representing African, European, Native American, and East Asian were selected from the Human Diversity Genome Project (HGDP) [42]. Each subpopulation (populations within each continent) was separately filtered for MAF >= 5% and for HWE p-value < 1e-8 using PLINK v1.9 [43]. Individuals with a relatedness degree, measured by the kinship coefficient using KING greater than 0.0884 were removed [44]. Admixed individuals in the reference dataset were removed, detected in a non-supervised ADMIXTURE analysis, with a threshold of 95% [45]. We balanced the reference population for ancestral representation, randomly removing samples from overrepresented populations. After filtering, we merged the reference dataset with the admixed dataset to obtain a list of overlapping SNPs used in LA estimation. LA inference was performed using G-Nomix [46]. The LA of each *APOE* allele was defined as African (AFR), European (EUR), Native American (NAM), or East Asian (EAS), considering a window of considering a window of 2 MB around the *APOE* locus (chr19:42,000,000-46,000,000). This window is the same previously used to fine map *APOE*-LA interactions in AD [26]. EAS ancestry was removed due to low frequency of admixed alleles. Global ancestry was estimated by modeling four ancestral populations (k = 4) with admixture in Structure version 2.3.3 (100,000 burn-ins, 200,000 iterations, LOCPRIOR = 0), alongside samples from the Human Genome Diversity Panel and the HapMap (Phase I) project, both publicly available [24, 29]. For analyses involving local *APOE* ancestry, admixed individuals presented combinations such as EUR/EUR, EUR/AFR, EUR/NAM, AFR/AFR, AFR/EUR, AFR/NAM, NAM/NAM, NAM/EUR, and NAM/AFR. After removing individuals with low confidence LA calls and/or EAS ancestry, the dataset contained 716 individuals from BAS and 952 individuals from SABE.

### Statistical Analysis

The sample collections were first characterized by sex, *APOE* genotype, and age (age at death for BAS; age at genetic material and data collection for SABE), with means and standard deviations presented in Table 1. Fisher’s exact tests were used to compare allele, genotype, and LA frequencies between BAS and SABE. Adherence to Hardy-Weinberg Equilibrium (HWE) was tested using Monte Carlo simulations with 10,000 permutations: the observed genotype counts were compared to the distribution of the genotype counts conditional on the observed allele counts, yielding a P-value for each genotype. The method for sampling genotype counts under HWE was the conventional method described in Guo and Thompson, 1992 . Since ancestry itself can cause deviations from HWE, an extended version of the simulations was also performed, accounting for LA. In these LA-information tests, the counts for each *APOE* + LA genotype combination were compared to their respective conditional distributions, assuming (A) fixed counts for each LA pair, (B) fixed counts for each *APOE* + LA allele combination, and (C) conditional independence between alleles (e.g., the expected proportion of ε4-EUR/ε3-AFR among EUR/AFR individuals is given by the product of the ε4 frequency in Europeans and the ε3 frequency in Africans). The Monte Carlo approach was implemented for all HWE tests, including *APOE* genotype frequency and *APOE* + LA genotype combinations, to account for low expected counts in certain low-frequency alleles and admixed combinations. Adjustment for multiple tests was applied to the results of each ancestry group, using the Westfall-Young method [47]. Furthermore, the distribution of LA relative to Global Ancestry (GA) was compared within each sample collection to assess potential selection or sampling bias. This analysis was performed using paired t-tests, comparing ancestry proportions within the *APOE* window to the ancestry proportions in the whole chromosome 19.

**Table 1.** Demographic, *APOE* genotype, and local ancestry (LA) distribution in the BAS and SABE sample collection.

|  | sample collection |  |  |
| --- | --- | --- | --- |
|  | BAS | SABE | P-value |
| **n** | 716 | 952 |  |
| **Female (%)** | 50.97 | 60.27 |  |
| **Mean age (sd)** | 72.7 (±12.5) | 72.37 (±9.48) | |
| **Genotypes** | ***APOE* Genotype frequency (Count)** |  |  |
| ε2/ε2 | 0.8 (6) | 0.1 (1) | 0.05 |
| ε2/ε3 | 9.2 (66) | 10.3 (98) | 0.51 |
| ε2/ε4 | 2.2 (16) | 2.2 (21) | 1.00 |
| ε3/ε3 | 60.5 (433) | 66.7 (635) | 0.01 |
| ε3/ε4 | 24.4 (175) | 18.4 (175) | 0.003 |
| ε4/ε4 | 2.8 (20) | 2.3 (22) | 0.50 |
| **Alles** | ***APOE* Allele frequency (Count)** |  |  |
| ε2 | 6.6 (94) | 6.4 (121) | 0.83 |
| ε3 | 77.3 (1107) | 81 (1543) | 0.01 |
| ε4 | 16.1 (231) | 12.6 (240) | 0.00 |
| **Ancestries** | ***APOE* Local Ancestry genotype Frequency (Count)** |  |  |
| **AFR/AFR** | 14.1 (101) | 7.4 (70) | <0.001 |
| **AFR/EUR** | 27.2 (195) | 17.5 (167) | <0.001 |
| **AFR/NAM** | 4.5 (32) | 2.7 (26) | 0.06 |
| **EUR/EUR** | 46.2 (331) | 62.9 (599) | <0.001 |
| **EUR/NAM** | 8 (57) | 8.7 (83) | 0.59 |
| **NAM/NAM** | 0 (0) | 0.7 (7) | 0.02 |
| **Ancestries** | ***APOE* Local Ancestry Allele Frequency (Count)** |  |  |
| **AFR** | 30 (429) | 17.5 (333) | <0.001 |
| **EUR** | 63.8 (914) | 76.1(1448) | <0.001 |
| **NAM** | 6.2 (89) | 6.5 (123) | 0.83 |

Summary of demographic characteristics, *APOE* genotypes, and local ancestry proportions in the BAS and SABE. Values are presented as percentages or mean ± standard deviation, as appropriate. Differences between sample collections were assessed using t-tests, with significant p-values ( $p < 0.05$ ) indicated.

## RESULTS

This study included 716 samples from the Brain Bank (BAS) and 952 from the SABE (see Table 1). The mean age was similar in both sample collections (72.7 ± 12.5 years in BAS and 72.37 ± 9.48 years in SABE). The proportion of females was 50.97% in BAS and 60.27% in SABE.

Regarding *APOE* genotypes, ε3/ε3 was the most frequent in both, being significantly higher in SABE (66.7%) than in BAS (60.5%) (p = 0.01). Correspondingly, the ε3 allele frequency was also higher in SABE (81.0%) than in BAS (77.3%) (p = 0.009). The ε2/ε2 genotype was significantly more prevalent in BAS (0.8%) than in SABE (0.1%) (p = 0.047), despite similar overall ε2 allele frequencies (6.6% and 6.4%, respectively; p = 0.831). The ε3/ε4 genotype was also significantly more frequent in BAS (24.4%) than in SABE (18.4%) (p=0.003).

LA analysis revealed a significantly higher proportion of African ancestry in the BAS sample collection (p < 0.001; Table 1). This difference was most evident in the proportion of haplotypes with AFR/AFR LA configurations, which accounted for 14.1% in BAS versus 7.1% in SABE (p < 0.001). Conversely, haplotypes with EUR/EUR LA configurations were significantly more prevalent in SABE (62.9%) than in BAS (46.2%) (p < 0.001). The proportion of ancestry heterozygotes (AFR/EUR admixture) was also higher in BAS (27.7%) than in SABE (17.5%) (p < 0.001).

When analyzing the combined *APOE* allele + LA haplotype frequencies (see Table 2), the frequency of the ε2-African allele (ε2 on an African LA background) was higher in the BAS (12.6%) than in the SABE (9.9%). Conversely, the ε2-European frequency was higher in SABE (6.1%) than in BAS (4.4%). The ε2-Native American (NAM) haplotype was not detected in either sample collections. There was no significant difference in the frequencies of the ε3-African (59.9% and 62.5%, in the BAS and SABE, respectively) and ε3-European (85.7% and 85.8%, respectively) haplotypes. Similar to the ε3 allele, the ε4-African haplotype showed the same proportions (27.5% in BAS vs. 27.6% in SABE), with no significant difference. However, the ε4-European haplotype was more prevalent in the BAS (10.0%) than in the SABE (8.1%). The ε3-Native American and ε4-NAM haplotypes also showed comparable proportions (75.0% and 24.0%, respectively). Importantly, none of the *APOE*+LA haplotype comparisons were statistically significant (p < 0.05) (See Supplementary Table S2).

**Table 2.**
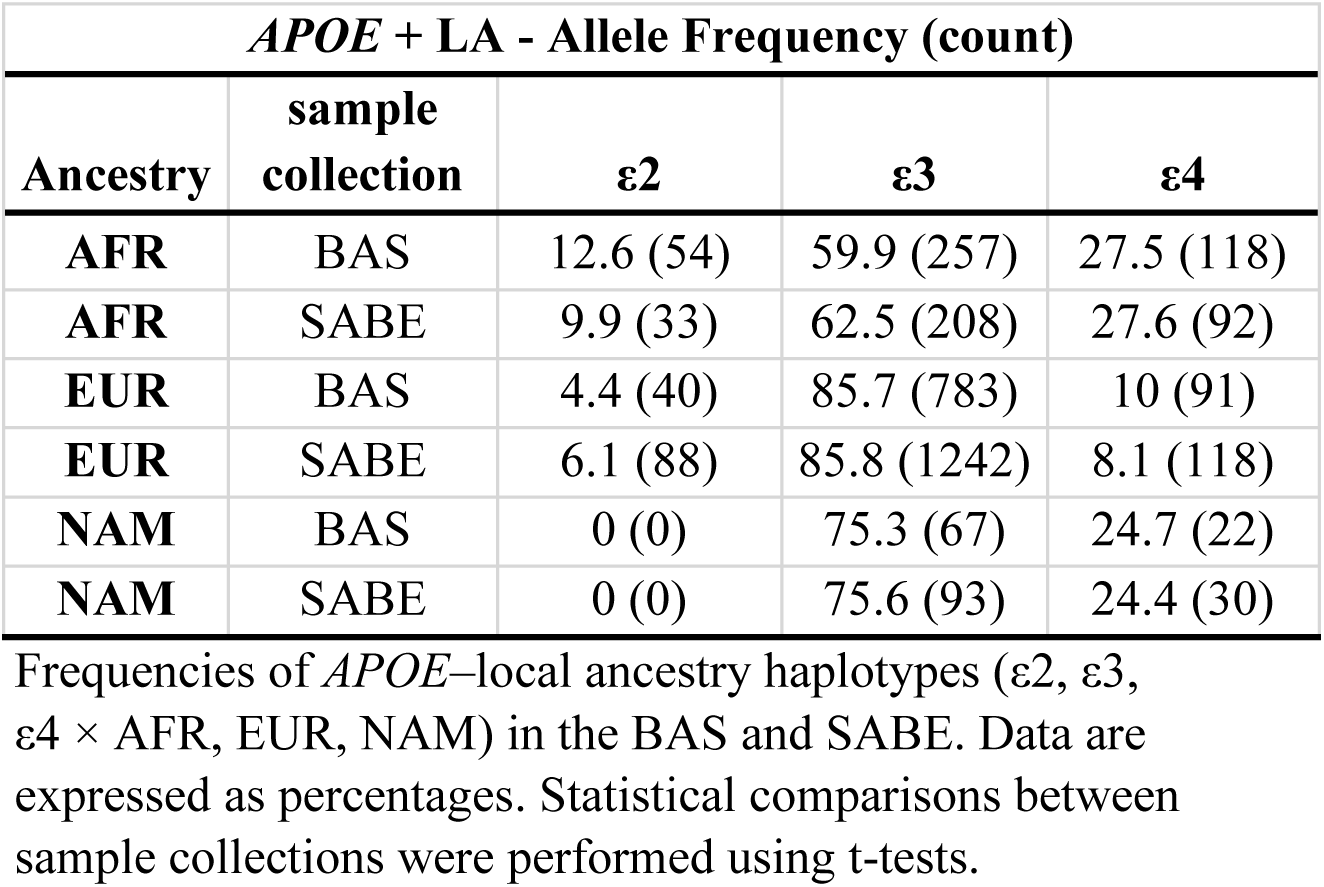
Haplotype frequencies combining *APOE* + LA in the BAS and SABE sample collection.

| ***APOE* + LA - Allele Frequency (count)** |  |  |  |  |
| --- | --- | --- | --- | --- |
| **Ancestry** | **sample collection** | **$\epsilon 2$** | **$\epsilon 3$** | **$\epsilon 4$** |
| **AFR** | BAS | 12.6 (54) | 59.9 (257) | 27.5 (118) |
| **AFR** | SABE | 9.9 (33) | 62.5 (208) | 27.6 (92) |
| **EUR** | BAS | 4.4 (40) | 85.7 (783) | 10 (91) |
| **EUR** | SABE | 6.1 (88) | 85.8 (1242) | 8.1 (118) |
| **NAM** | BAS | 0 (0) | 75.3 (67) | 24.7 (22) |
| **NAM** | SABE | 0 (0) | 75.6 (93) | 24.4 (30) |

Frequencies of *APOE*–local ancestry haplotypes ( $\epsilon 2$ , $\epsilon 3$ , $\epsilon 4 \times$ AFR, EUR, NAM) in the BAS and SABE. Data are expressed as percentages. Statistical comparisons between sample collections were performed using t-tests.

We first tested *APOE* genotype frequencies for deviations from HWE using Monte Carlo simulations due to low sample counts for some alleles. No significant HWE deviation was observed in either sample collections, except the underrepresented ε3/ε4 genotype in SABE (observed = 175, expected =194;5; p = 0.0083; adjusted p =0.032) (See Supplementary Tables S3 and S4).

However, given that LA can reflect sub-population structure, we next tested HWE by accounting for LA at the *APOE* locus, evaluating over- or underrepresentation in LA genotypes irrespective of the *APOE* allele. We observed a significant HWE deviation at the LA genotype level in both sample collections, driven by a consistent pattern across the major ancestral groups. Specifically, both BAS and SABE showed a significant overrepresentation of LA-homozygotes: AFR/AFR (BAS: observed=101, expected=64.1; SABE: observed=70, expected=29.1) and EUR/EUR (BAS: observed=331, expected=291.6; SABE: observed=599, expected=550.6). LA-heterozygotes AFR/EUR were significantly underrepresented in both sample collections (BAS: observed=195, expected=274.5; SABE: observed=167, expected=253.2) (See Supplementary Tables S5 and S6). For all these major LA genotypes, the deviations were highly significant (adjusted p < 2 x 10^-4). Deviations in LA genotypes involving NAM were not significant after multiple test correction, likely due to the smaller sample size.

We compared the distribution of LA relative to GA within each sample collection. The BAS samples exhibited significant differences: specifically, a higher proportion of European LA (1.3%; 95% CI: 0.13 to 2.57%; p = 0.031) and a lower proportion of Native American LA (−0.95%; 95% CI: −1.71 to −0.19%; p = 0.014). In contrast, the SABE did not show any statistically significant differences in the same comparisons (p > 0.05) (See Supplementary Table S1).

Building upon the observed LA deviations, we tested a more general genotypic model, which treats LA groups as exogenous (fixed) and accounts for differences in allele frequency across LA groups. The results revealed a significant overrepresentation (excess) of the ε2 AFR/ε2 AFR genotype combination in the BAS (observed= 5, expected= 1.6; p = 0.0196; adjusted p = 0.093) (See Figure 1, Supplementary Tables S7 and S8). In contrast, HWE deviations in the SABE were widespread, primarily showing overrepresentations in ε3-AFR/ε3-EUR (observed=107, expected=89.5; p=0.0003; adjusted p = 0.0014), ε4-AFR/ε4-AFR (observed=11, expected=5.3; p=0.009; adjusted p = 0.033) and ε2-AFR/ε3-NAM (observed=5, expected=1.9; p=0.0424; adjusted p = 0.16).

**Figure 1.**
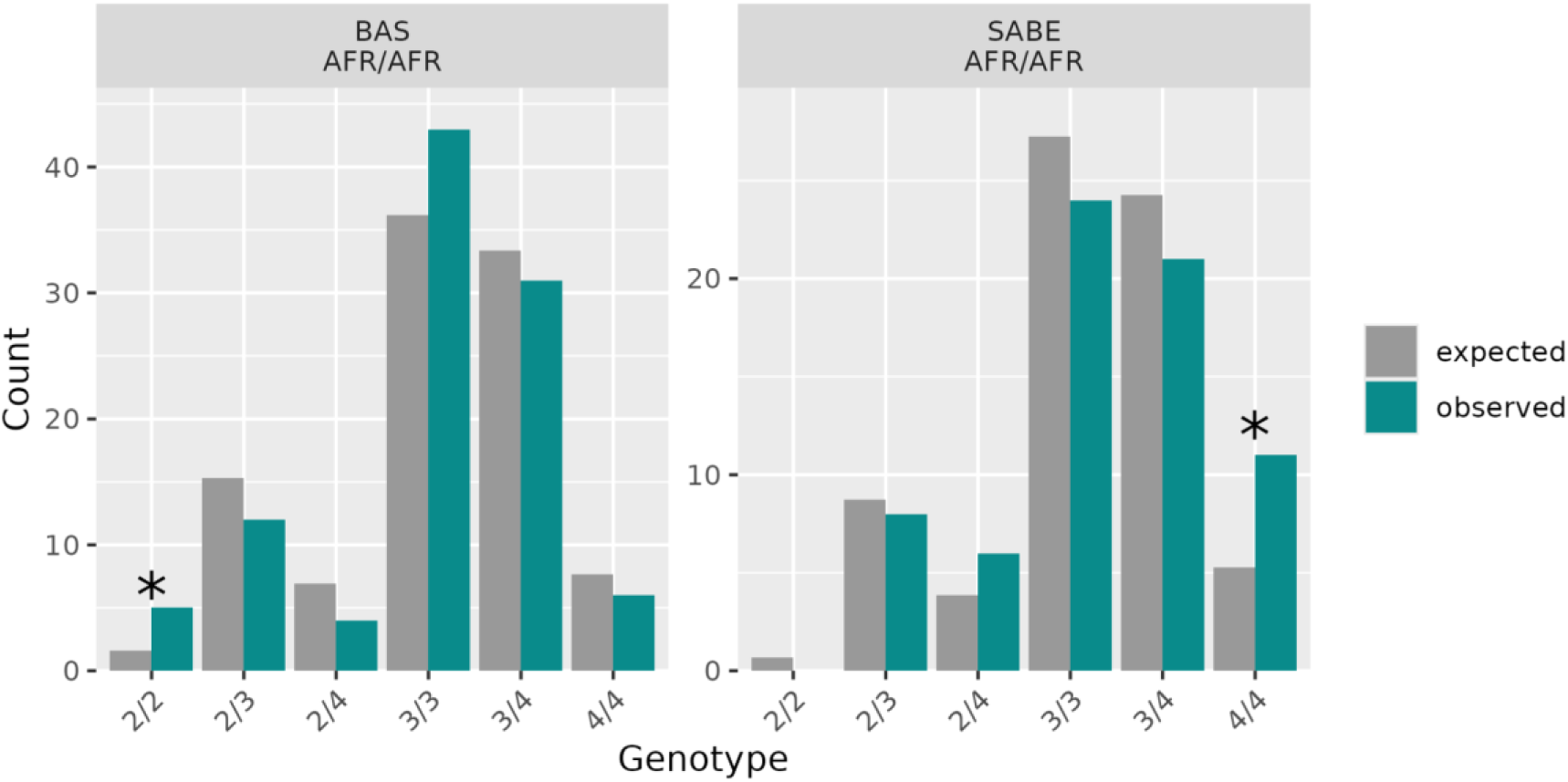
Expected vs. observed genotype counts for AFR/AFR individuals. Expected counts derived via Monte Carlo simulation. Asterisk indicates adjusted p < 0.10.

**Figure 2.**
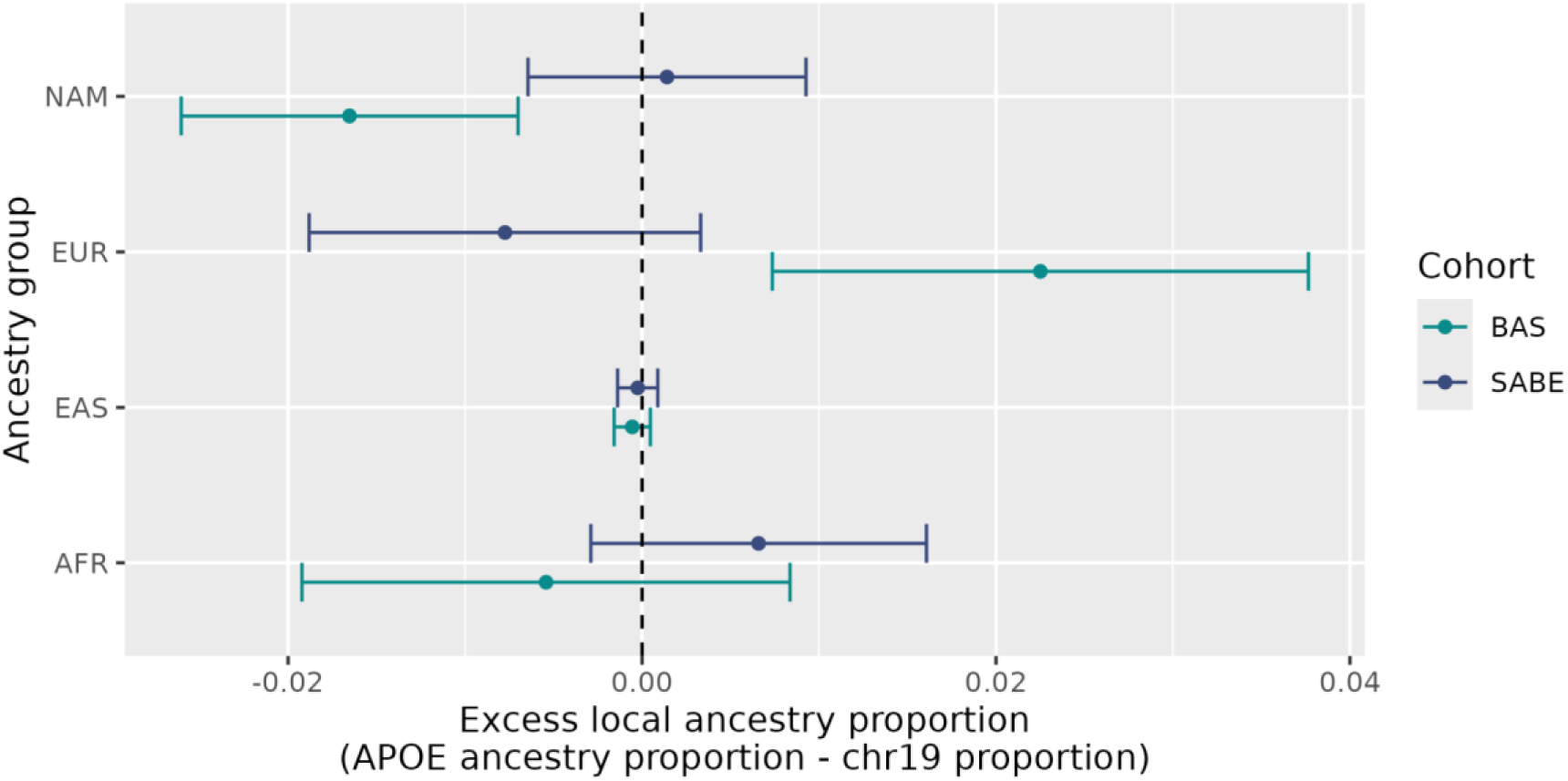
Local ancestry enrichment at the *APOE* locus. Differences between ancestry proportions at the *APOE* locus and the entire chromosome 19. Error bars indicate 95% confidence intervals.

## DISCUSSION

*APOE* genotype frequencies showed no significant deviation from HWE in either sample collection, suggesting the locus itself is not under strong assortative mating at a panmictic level. HWE serves as a null model; a lack of deviation implies that allele frequencies remain stable over generations in the absence of targeted selective pressures [38–40]. However, LA analysis revealed highly significant deviations from HWE, characterized by a pervasive excess of LA homozygotes and a deficit of heterozygotes across major ancestral groups. This pattern strongly suggests assortative mating with regard to ancestry, likely mediated by historical and socioeconomic geographical factors in São Paulo. The deviation from HWE for LA, therefore, reflects underlying population structure rather than targeted selection. Recognizing this, the incorporation of LA into subsequent genotypic models to avoid stratification biases is highly recommended [38].

We observed significant differences in genotype composition and ancestry between the BAS and SABE sample collections, which can lead to a crucial distinction for interpreting risk variation [24, 26, 28, 29, 48]. Overall, the post-mortem BAS sample exhibits greater African ancestry and higher genetic diversity, driven by a higher proportion of *ε3/ε4* heterozygotes. In contrast, the SABE is more European and homogeneous, with a predominance of the *ε3/ε3* genotype. Despite its greater African ancestry, the BAS showed a significant excess of European LA compared to its GA, accompanied by a deficit of Native American LA. This localized asymmetry suggests complex population dynamics and potential differential mortality pressures operating within specific ancestral backgrounds [6, 12, 13].

The excess of the *ε4*-European haplotype in the post-mortem BAS sample collection aligns with previous literature linking this specific combination to increased AD risk and accelerated mortality. These findings reinforce the idea that LA modulates clinical risks, where the *ε4* allele exerts its full deleterious impact primarily when embedded within a European genetic background.

Most importantly, our study extends this ancestry-driven modulation to the opposite end of the risk spectrum, raising crucial questions about the assumed universality of the protective *ε2* allele.Detecting evolutionary selection signatures in rare variants under admixed backgrounds is statistically challenging due to low expected genotype counts. To robustly test our generalized genotypic model, we employed Monte Carlo simulations with 10,000 permutations, ensuring accurate HWE deviation estimates irrespective of sparse sampling, fortified by Westfall-Young step-down adjustments for multiple comparisons. Using this rigorous framework, a central finding of our study was the significant HWE deviations observed when treating LA groups under a general genotypic model: an excess of the *ε2*AFR/*ε2*AFR genotype combination in the deceased sample collection (BAS), contrasted with an overrepresentation of *ε4*AFR/*ε4*AFR in the living sample collection (SABE). We argue that these mirrored deviations indicate that an African LA background acts as a genetic buffer that attenuates the phenotypic extreme effects of *APOE* alleles in AD pathology and longevity. In the case of *ε4*, this buffering effect mitigates its deleterious impact, enabling African-ancestry *ε4*/*ε4* carriers to reach advanced ages and accumulate in the living Sample collection (SABE).

Conversely, this same buffering mechanism appears to diminish the exceptional longevity advantage typically conferred by the *ε2* allele in European-centric studies. Because *ε2*AFR/*ε2*AFR individuals do not experience a distinct survival advantage over the rest of the population, they accumulate expectedly within the naturally deceased BAS sample collection. Therefore, these signals indicate that the APOE *ε2* allele may not behave as a universal longevity factor across all human populations. Instead, its optimal protective mechanisms may be highly contingent on specific genomic contexts, such as a European LA background, highlighting the need to reevaluate standard biomarkers within highly admixed, non-European populations.

Our findings provide a critical new lens through which to interpret previous literature on the *APOE ε2* allele. While GWAS and epidemiological studies have robustly associated *ε2* with exceptional longevity and protection against AD [1, 2, 6, 12–14, 20], these prevailing conclusions have been overwhelmingly drawn from populations of European descent [49–51]. Furthermore, because local ancestry inference is methodologically demanding and the *ε2* allele is globally rare, most prior studies in admixed populations have either relied solely on Global Ancestry (GA) [6] or pooled *ε2* genotypes (combining heterozygotes and rare homozygotes) to achieve statistical power, as highlighted by Suri et al. (2013) [31]. This methodological constraint inherently masks sub-population genomic dynamics. By disentangling the *APOE* locus using high-resolution LA inference, our results suggest that the historically reported heterogeneity in *ε2* frequencies and its protective effects across diverse populations may not merely be statistical noise or environmental variance, but rather a biological reality shaped by the local genomic background. Therefore, the lack of a universal survival advantage for the *ε2* allele observed in our African LA background challenges the “one-size-fits-all” assumption of *APOE* biomarkers, emphasizing that the evolutionary and clinical implications of “the forgotten allele” are deeply context-dependent.

A key strength of our study is the novel approach of contrasting a post-mortem (BAS) with a living (SABE) sample collection to capture survival dynamics in a highly admixed population. However, we acknowledge that these collections were acquired in distinct contexts. The BAS is a community-dwelling biobank of individuals deceased from natural causes, often characterized by a high prevalence of vascular endophenotypes. In contrast, the SABE is a census-based longitudinal study comprising living, generally healthier older adults [29, 36]. While these differing recruitment settings might partially explain some of the general ancestry deviations observed, there is no established evidence that these demographic and methodological parameters induce targeted selection bias specifically for *APOE* genotypes. Crucially, our HWE analyses support this premise; the models do not point to direct, unmeasured selection forces acting on the *APOE* locus that cannot be explained by geographical conditions and underlying population structure.

A notable limitation of this study is the lack of adjustment for educational attainment, which is a critical factor for cognitive reserve and dementia outcomes. Nevertheless, because our primary focus is strictly on longevity, assessed through mortality versus survival proxies, rather than cognitive performance trajectories, this limitation primarily restricts direct cognitive comparisons between the cohorts. Consequently, we deliberately refrained from making cross-cohort inferences regarding cognition, ensuring that our conclusions regarding the intersection of *APOE*, local ancestry, and longevity remain methodologically reliable.

## Supporting information

all suplementary data

## ACKNOWLEDGMENTS

We thank our research participants and family members for their contribution to this study. Funding was provided by Fundação de Amparo à Pesquisa do Estado de São Paulo (FAPESP 2025/00999-5, 2022/03102-8, CEPID 2013/08028-1, 2014/50649-6, INCT 2014/50931-3), Conselho Nacional de Desenvolvimento Científico e Tecnológico (CNPq INCT 465355/2014-5, 304746/2022-3,), Instituto Serrapilheira (Call 6), Alzheimer’s Association (24CBIDR-1185483, 23AARG-1026658)

---

**Table S1:**
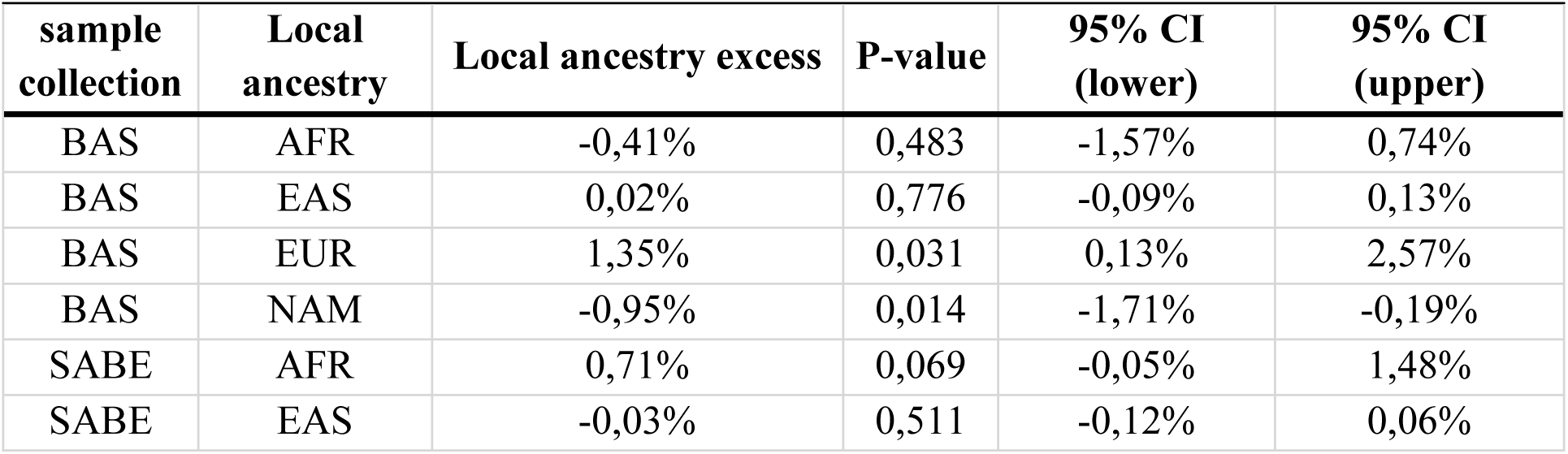
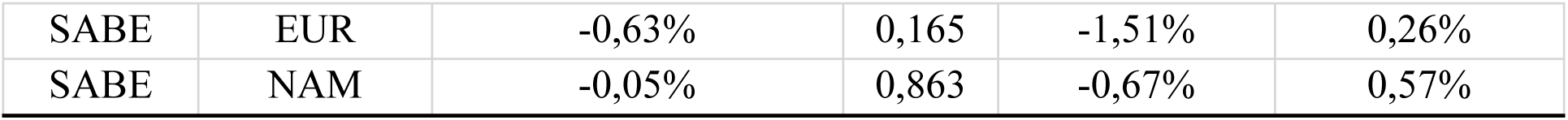
Comparison of Local Ancestry (LA) Proportion Relative to Global Ancestry (GA) in the BAS and SABE. This table details the mean percentage difference between LA and GA proportions for African (AFR), Native American (NAM), and European (EUR) ancestries within the BAS and SABE. The percentage difference is followed by p-value and the corresponding 95% Confidence Interval (95% CI, upper and lower bounds), demonstrating internal ancestry deviation in each sample collection.

**Table S2:**
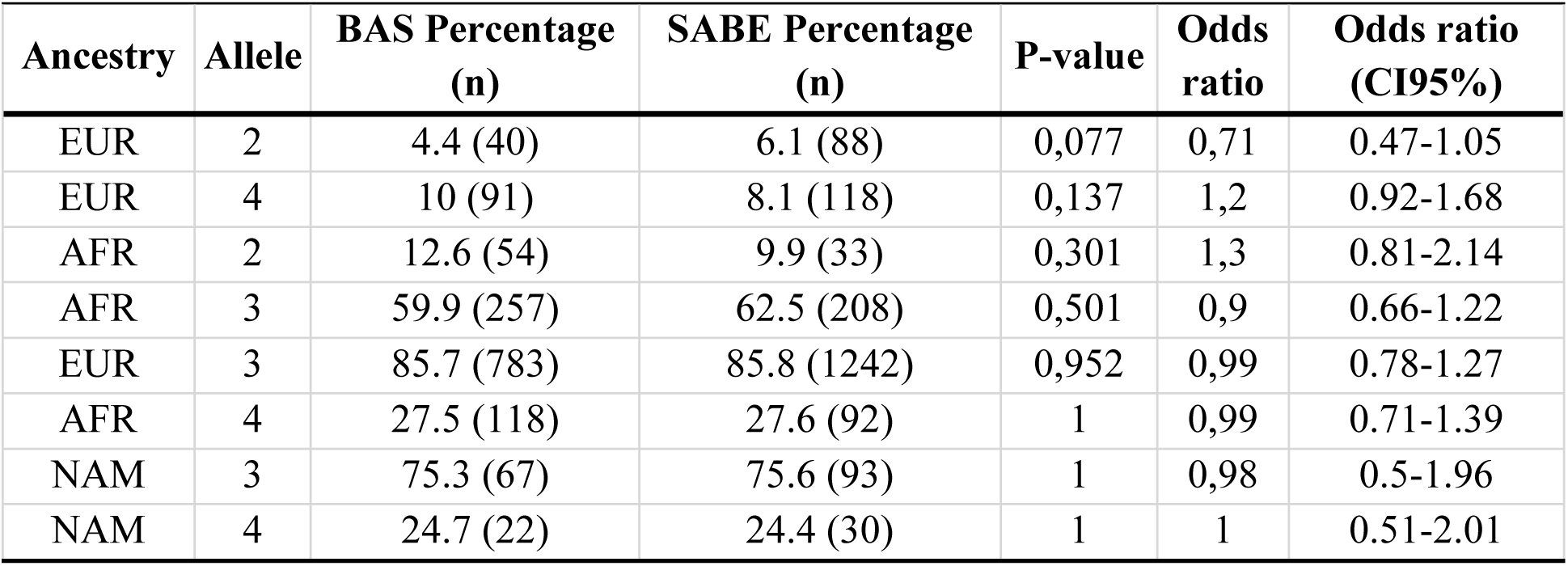
Comparison of *APOE* Allele Frequencies Stratified by Local Ancestry (LA) and Differential Association with sample collection Status.This table presents the observed counts and frequencies (%) of the *APOE* alleles (ε2, ε3, ε4) combined with their corresponding LA (African, European, Native American) in the post-mortem BAS and SABE). The differential frequency of each *APOE* + LA haplotype between the is quantified by the Odds Ratio (OR), its 95% Confidence Interval (95% CI), and the corresponding p-value.

**Table S3:**
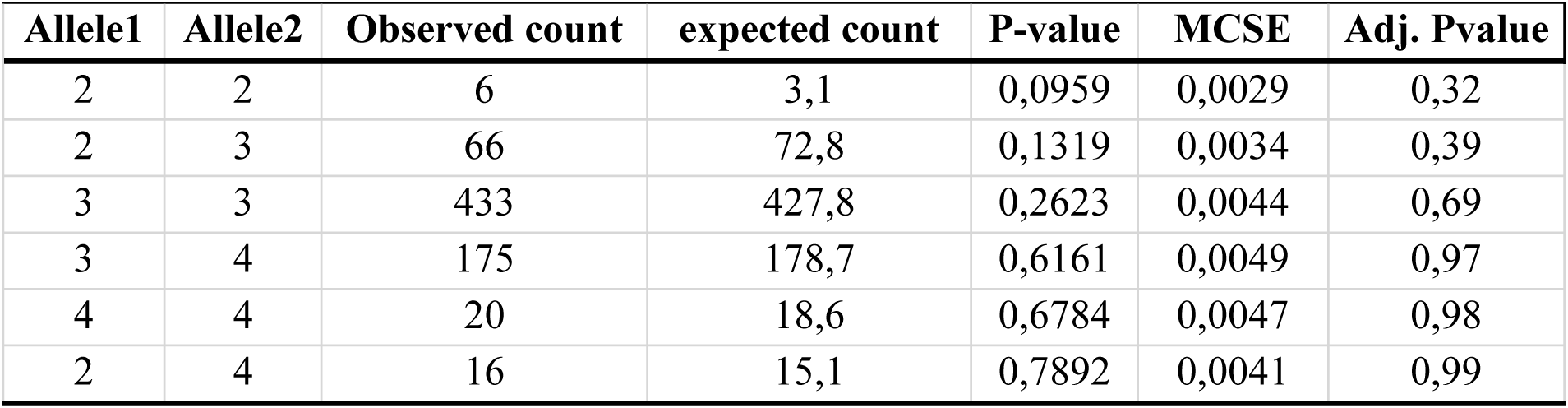
Hardy-Weinberg Equilibrium (HWE) Analysis for *APOE* Genotype Frequencies in the BAS sample collection. The table presents the comprehensive results of the HWE test for all *APOE* genotype combinations in the BAS sample collection. Shown are the observed counts, expected counts, p-values, Monte Carlo Standard Errors (MCSE), and adjusted p-values.

**Table S4:**
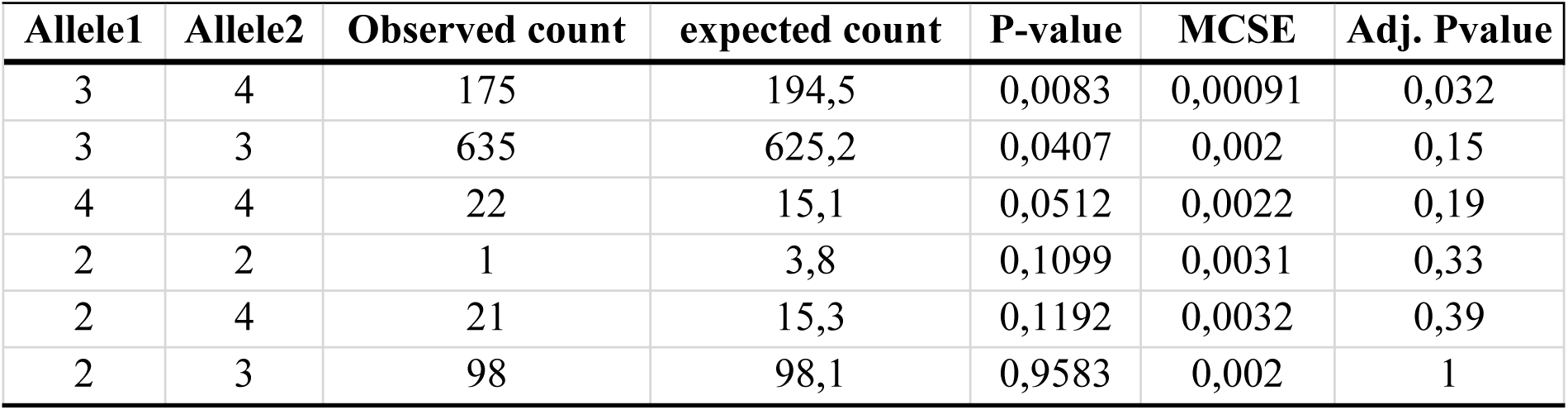
Hardy-Weinberg Equilibrium (HWE) Analysis for *APOE* Genotype Frequencies in the SABE. . The table presents the comprehensive results of the HWE test for all *APOE* genotype combinations in the SABE. Shown are the observed counts, expected counts, p-values, Monte Carlo Standard Errors (MCSE), and adjusted p-values.

**Table S5:**
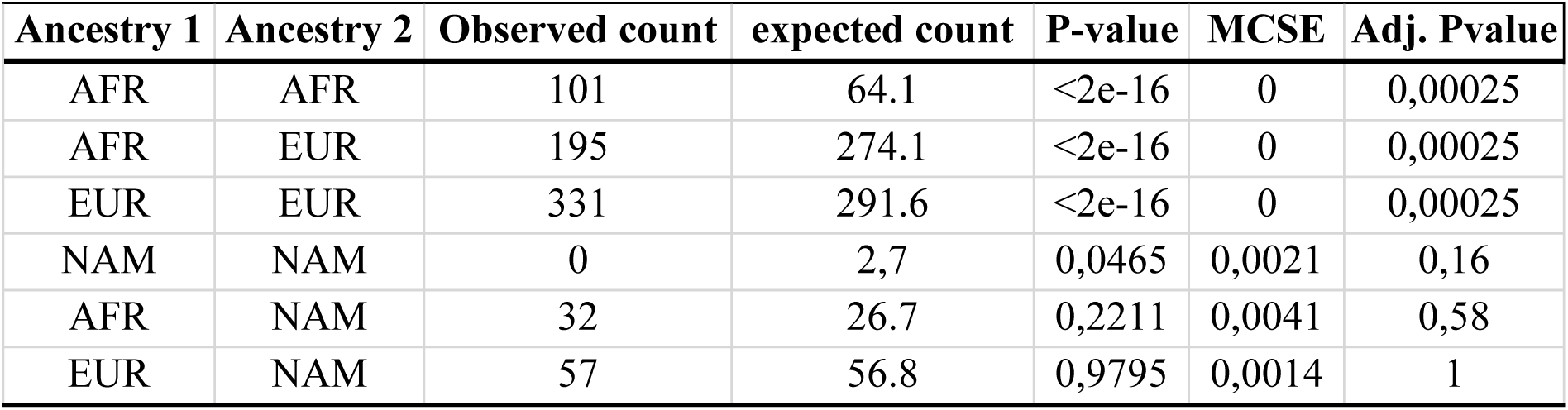
Hardy-Weinberg Equilibrium (HWE) Analysis for Local Ancestry (LA) Genotype Frequencies in the BAS. The table presents the comprehensive results of the HWE test for all LA genotype combinations in the BAS. Shown are the observed counts, expected counts, p-values, Monte Carlo Standard Errors (MCSE), and adjusted p-values.

**Table S6:**
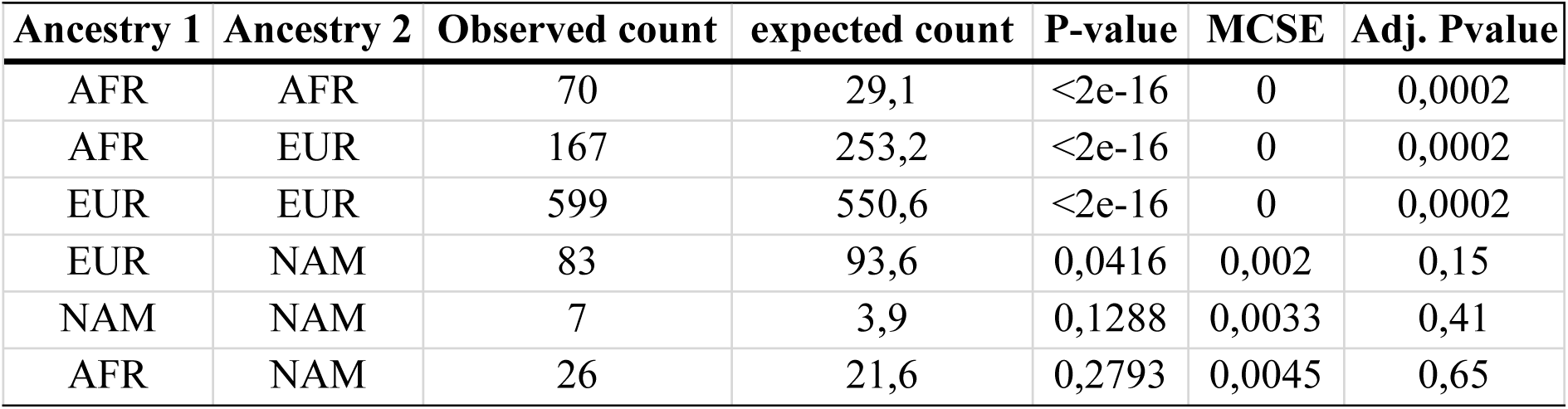
Hardy-Weinberg Equilibrium (HWE) Analysis for Local Ancestry (LA) Genotype Frequencies in the SABE. The table presents the comprehensive results of the HWE test for all LA genotype combinations in the SABE. Shown are the observed counts, expected counts, p-values, Monte Carlo Standard Errors (MCSE), and adjusted p-values.

**Table S7:**
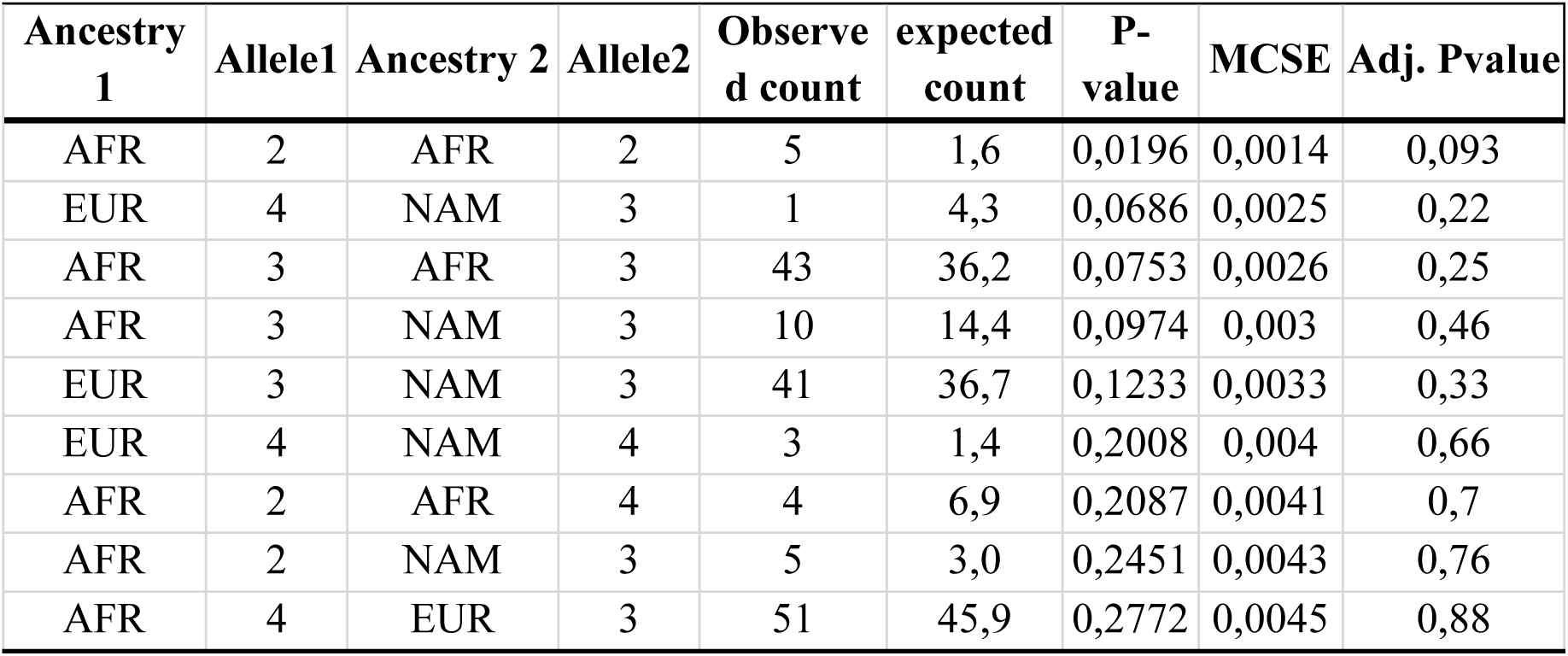
Hardy-Weinberg Equilibrium (HWE) Analysis for *APOE* + Local Ancestry (LA) Genotypes in the BAS. The table presents the comprehensive results of the HWE test for all *APOE* genotype combinations stratified by LA background in the BAS. Shown are the observed counts, expected counts, p-values, Monte Carlo Standard Errors (MCSE), and adjusted p-values.

**Table S8:**
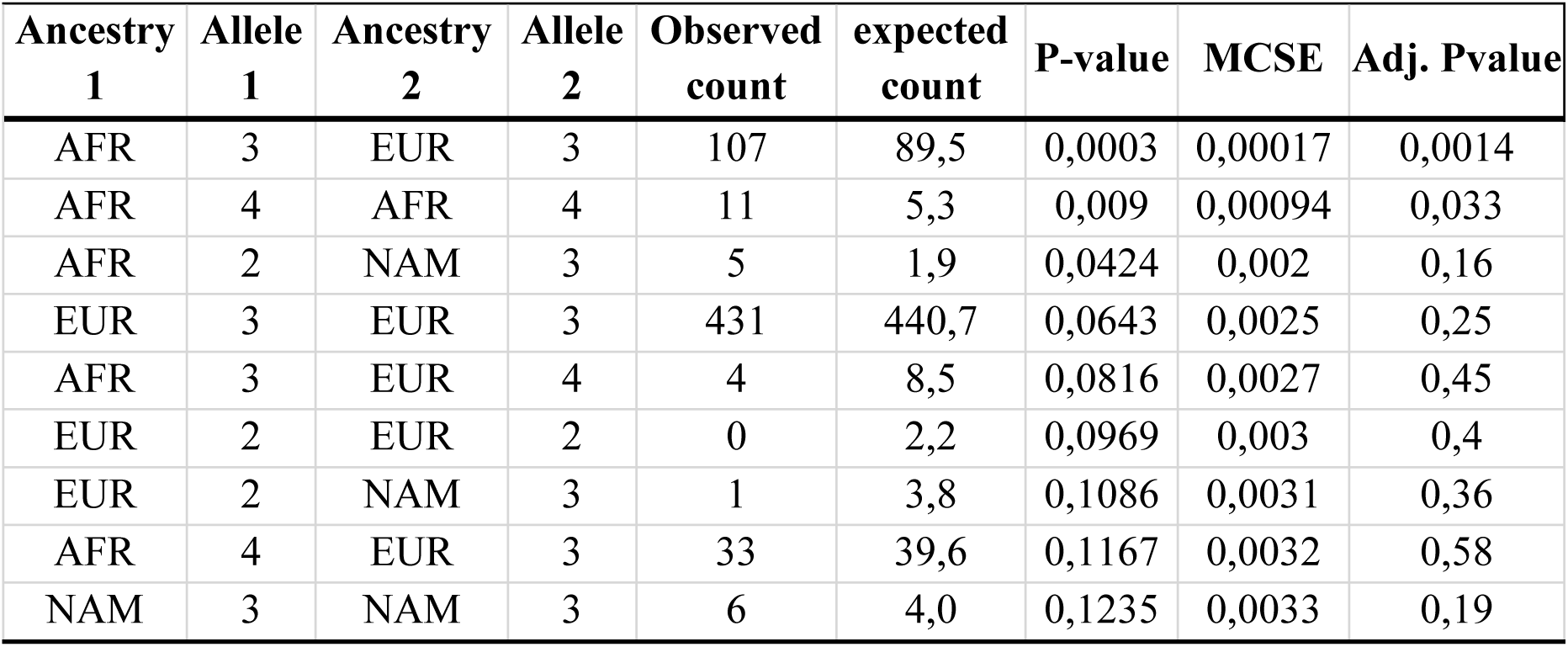
Hardy-Weinberg Equilibrium (HWE) Analysis for *APOE* + Local Ancestry (LA) Genotypes in the BAS. The table presents the comprehensive results of the HWE test for all *APOE* genotype combinations stratified by LA background in the BAS. Shown are the observed counts, expected counts, p-values, Monte Carlo Standard Errors (MCSE), and adjusted p-values.

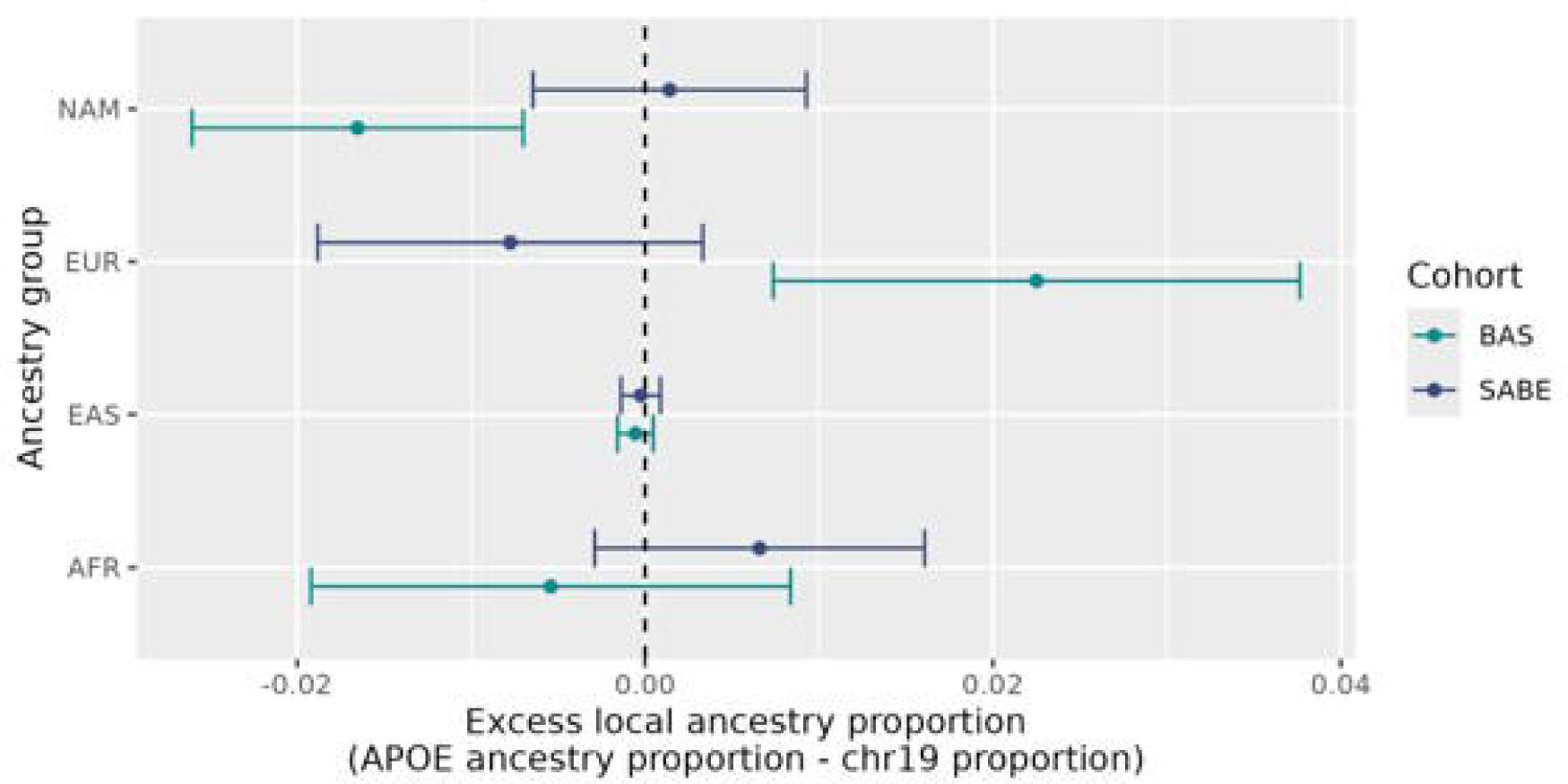

