## Supplementary material for "Is *APOE ε2* always a protective allele? Deviations in Hardy-Weinberg equilibrium in admixed Brazilian elderly individuals": all suplementary data

**SUPPLEMENTARY TABLES**

| sample collection | Local ancestry | Local ancestry excess | P-value | 95% CI (lower) | 95% CI (upper) |
|---|---|---|---|---|---|
| BAS | AFR | -0,41% | 0,483 | -1,57% | 0,74% |
| BAS | EAS | 0,02% | 0,776 | -0,09% | 0,13% |
| BAS | EUR | 1,35% | 0,031 | 0,13% | 2,57% |
| BAS | NAM | -0,95% | 0,014 | -1,71% | -0,19% |
| SABE | AFR | 0,71% | 0,069 | -0,05% | 1,48% |
| SABE | EAS | -0,03% | 0,511 | -0,12% | 0,06% |
| SABE | EUR | -0,63% | 0,165 | -1,51% | 0,26% |
| SABE | NAM | -0,05% | 0,863 | -0,67% | 0,57% |

**Table S1: Comparison of Local Ancestry (LA) Proportion Relative to Global Ancestry (GA) in the BAS and SABE.** This table details the mean percentage difference between LA and GA proportions for African (AFR), Native American (NAM), and European (EUR) ancestries within the BAS and SABE. The percentage difference is followed by p-value and the corresponding 95% Confidence Interval (95% CI, upper and lower bounds), demonstrating internal ancestry deviation in each sample collection.

| Ancestry | Allele | BAS Percentage (n) | SABE Percentage (n) | P-value | Odds ratio | Odds ratio (CI95%) |
|---|---|---|---|---|---|---|
| EUR | 2 | 4.4 (40) | 6.1 (88) | 0,077 | 0,71 | 0.47-1.05 |
| EUR | 4 | 10 (91) | 8.1 (118) | 0,137 | 1,2 | 0.92-1.68 |
| AFR | 2 | 12.6 (54) | 9.9 (33) | 0,301 | 1,3 | 0.81-2.14 |
| AFR | 3 | 59.9 (257) | 62.5 (208) | 0,501 | 0,9 | 0.66-1.22 |
| EUR | 3 | 85.7 (783) | 85.8 (1242) | 0,952 | 0,99 | 0.78-1.27 |
| AFR | 4 | 27.5 (118) | 27.6 (92) | 1 | 0,99 | 0.71-1.39 |
| NAM | 3 | 75.3 (67) | 75.6 (93) | 1 | 0,98 | 0.5-1.96 |
| NAM | 4 | 24.7 (22) | 24.4 (30) | 1 | 1 | 0.51-2.01 |

**Table S2: Comparison of *APOE* Allele Frequencies Stratified by Local Ancestry (LA) and Differential Association with sample collection Status.** This table presents the observed counts and frequencies (%) of the *APOE* alleles (ε2, ε3, ε4) combined with their corresponding LA (African, European, Native American) in the post-mortem BAS and SABE). The differential frequency of each *APOE* + LA haplotype between the is quantified by the Odds Ratio (OR), its 95% Confidence Interval (95% CI), and the corresponding p-value.

| Allele1 | Allele2 | Observed count | expected count | P-value | MCSE | Adj. Pvalue |
|---|---|---|---|---|---|---|
| 2 | 2 | 6 | 3,1 | 0,0959 | 0,0029 | 0,32 |
| 2 | 3 | 66 | 72,8 | 0,1319 | 0,0034 | 0,39 |
| 3 | 3 | 433 | 427,8 | 0,2623 | 0,0044 | 0,69 |
| 3 | 4 | 175 | 178,7 | 0,6161 | 0,0049 | 0,97 |
| 4 | 4 | 20 | 18,6 | 0,6784 | 0,0047 | 0,98 |
| 2 | 4 | 16 | 15,1 | 0,7892 | 0,0041 | 0,99 |

**Table S3: Hardy-Weinberg Equilibrium (HWE) Analysis for *APOE* Genotype Frequencies in the BAS sample collection.** The table presents the comprehensive results of the HWE test for all *APOE* genotype combinations in the BAS sample collection. Shown are the observed counts, expected counts, p-values, Monte Carlo Standard Errors (MCSE), and adjusted p-values.

| Allele1 | Allele2 | Observed count | expected count | P-value | MCSE | Adj. Pvalue |
|---|---|---|---|---|---|---|
| 3 | 4 | 175 | 194,5 | 0,0083 | 0,00091 | 0,032 |
| 3 | 3 | 635 | 625,2 | 0,0407 | 0,002 | 0,15 |
| 4 | 4 | 22 | 15,1 | 0,0512 | 0,0022 | 0,19 |
| 2 | 2 | 1 | 3,8 | 0,1099 | 0,0031 | 0,33 |
| 2 | 4 | 21 | 15,3 | 0,1192 | 0,0032 | 0,39 |
| 2 | 3 | 98 | 98,1 | 0,9583 | 0,002 | 1 |

**Table S4: Hardy-Weinberg Equilibrium (HWE) Analysis for *APOE* Genotype Frequencies in the SABE**. The table presents the comprehensive results of the HWE test for all *APOE* genotype combinations in the SABE. Shown are the observed counts, expected counts, p-values, Monte Carlo Standard Errors (MCSE), and adjusted p-values.

| Ancestry 1 | Ancestry 2 | Observed count | expected count | P-value | MCSE | Adj. Pvalue |
|---|---|---|---|---|---|---|
| AFR | AFR | 101 | 64.1 | <2e-16 | 0 | 0,00025 |
| AFR | EUR | 195 | 274.1 | <2e-16 | 0 | 0,00025 |
| EUR | EUR | 331 | 291.6 | <2e-16 | 0 | 0,00025 |
| NAM | NAM | 0 | 2,7 | 0,0465 | 0,0021 | 0,16 |
| AFR | NAM | 32 | 26.7 | 0,2211 | 0,0041 | 0,58 |
| EUR | NAM | 57 | 56.8 | 0,9795 | 0,0014 | 1 |

**Table S5: Hardy-Weinberg Equilibrium (HWE) Analysis for Local Ancestry (LA) Genotype Frequencies in the BAS.** The table presents the comprehensive results of the HWE test for all LA genotype combinations in the BAS. Shown are the observed counts, expected counts, p-values, Monte Carlo Standard Errors (MCSE), and adjusted p-values.

| Ancestry 1 | Ancestry 2 | Observed count | expected count | P-value | MCSE | Adj. Pvalue |
|---|---|---|---|---|---|---|
| AFR | AFR | 70 | 29,1 | <2e-16 | 0 | 0,0002 |
| AFR | EUR | 167 | 253,2 | <2e-16 | 0 | 0,0002 |
| EUR | EUR | 599 | 550,6 | <2e-16 | 0 | 0,0002 |
| EUR | NAM | 83 | 93,6 | 0,0416 | 0,002 | 0,15 |
| NAM | NAM | 7 | 3,9 | 0,1288 | 0,0033 | 0,41 |
| AFR | NAM | 26 | 21,6 | 0,2793 | 0,0045 | 0,65 |

**Table S6: Hardy-Weinberg Equilibrium (HWE) Analysis for Local Ancestry (LA) Genotype Frequencies in the SABE.** The table presents the comprehensive results of the HWE test for all LA genotype combinations in the SABE. Shown are the observed counts, expected counts, p-values, Monte Carlo Standard Errors (MCSE), and adjusted p-values.

| Ancestry 1 | Allele1 | Ancestry 2 | Allele2 | Observed count | expected count | P-value | MCSE | Adj. Pvalue |
|---|---|---|---|---|---|---|---|---|
| AFR | 2 | AFR | 2 | 5 | 1,6 | 0,0196 | 0,0014 | 0,093 |
| EUR | 4 | NAM | 3 | 1 | 4,3 | 0,0686 | 0,0025 | 0,22 |
| AFR | 3 | AFR | 3 | 43 | 36,2 | 0,0753 | 0,0026 | 0,25 |
| AFR | 3 | NAM | 3 | 10 | 14,4 | 0,0974 | 0,003 | 0,46 |
| EUR | 3 | NAM | 3 | 41 | 36,7 | 0,1233 | 0,0033 | 0,33 |
| EUR | 4 | NAM | 4 | 3 | 1,4 | 0,2008 | 0,004 | 0,66 |
| AFR | 2 | AFR | 4 | 4 | 6,9 | 0,2087 | 0,0041 | 0,7 |
| AFR | 2 | NAM | 3 | 5 | 3,0 | 0,2451 | 0,0043 | 0,76 |
| AFR | 4 | EUR | 3 | 51 | 45,9 | 0,2772 | 0,0045 | 0,88 |

**Table S7: Hardy-Weinberg Equilibrium (HWE) Analysis for *APOE* + Local Ancestry (LA) Genotypes in the BAS.** The table presents the comprehensive results of the HWE test for all *APOE* genotype combinations stratified by LA background in the BAS. Shown are the observed counts, expected counts, p-values, Monte Carlo Standard Errors (MCSE), and adjusted p-values.

| Ancestry 1 | Allele 1 | Ancestry 2 | Allele 2 | Observed count | expected count | P-value | MCSE | Adj. Pvalue |
|---|---|---|---|---|---|---|---|---|
| AFR | 3 | EUR | 3 | 107 | 89,5 | 0,0003 | 0,00017 | 0,0014 |
| AFR | 4 | AFR | 4 | 11 | 5,3 | 0,009 | 0,00094 | 0,033 |
| AFR | 2 | NAM | 3 | 5 | 1,9 | 0,0424 | 0,002 | 0,16 |
| EUR | 3 | EUR | 3 | 431 | 440,7 | 0,0643 | 0,0025 | 0,25 |
| AFR | 3 | EUR | 4 | 4 | 8,5 | 0,0816 | 0,0027 | 0,45 |
| EUR | 2 | EUR | 2 | 0 | 2,2 | 0,0969 | 0,003 | 0,4 |
| EUR | 2 | NAM | 3 | 1 | 3,8 | 0,1086 | 0,0031 | 0,36 |
| AFR | 4 | EUR | 3 | 33 | 39,6 | 0,1167 | 0,0032 | 0,58 |
| NAM | 3 | NAM | 3 | 6 | 4,0 | 0,1235 | 0,0033 | 0,19 |

**Table S8: Hardy-Weinberg Equilibrium (HWE) Analysis for *APOE* + Local Ancestry (LA) Genotypes in the BAS.** The table presents the comprehensive results of the HWE test for all *APOE* genotype combinations stratified by LA background in the BAS. Shown are the observed counts, expected counts, p-values, Monte Carlo Standard Errors (MCSE), and adjusted p-values.
